# Effect of READ1 on latent profiles of reading disability and comorbid attention and language learning disability subtypes

**DOI:** 10.1101/589226

**Authors:** Miao Li, Dongnhu T. Truong, Mellissa DeMille, Jeffrey G. Malins, Maureen W. Lovett, Joan Bosson-Heenan, Jeffrey R. Gruen, on behalf of the Genes, Reading, and Dyslexia Study, Jan C. Frijters

**Author notes:** Corresponding Author Jeffrey R. Gruen, 464 Congress Avenue, New Haven, CT 06520. Authors contributed equally to this work (Miao Li, Dongnhu T. Truong).

## Abstract

Recent studies of co-occurring reading disability (RD) and attention deficit/hyperactivity disorder (ADHD), and co-occurring RD and language learning disability (LLD), support a core disability plus co-occurrence model focused on language and attention. Genetic factors have been associated with poor reading performance. However, little is known about whether different genetic variants independently contribute to RD co-occurrence subtypes. We aimed to identify subgroups of struggling readers using a latent profile analysis (LPA) in a sample of 1,432 Hispanic American and African American youth. RD classes were then tested for association with variants of READ1, a regulatory element within the candidate RD risk gene, *DCDC2*. Six groups were identified in the LPA using RD designation as a known-class variable. The three RD classes identified groups of subjects with neurocognitive profiles representing RD+ADHD, specific phonological deficit RD, and RD+LLD. Genetic associations across RD subtypes were investigated against functional groupings of READ1. The RU1-1 group of READ1 alleles was associated with RD cases that were marked by deficits in both processing speed and attention (RD + ADHD). The *DCDC2* microdeletion that encompasses READ1 was associated with RD cases showing a phonological deficit RD profile. These findings provide evidence for differential genetic contribution to RD subtypes, and that previously implicated genetic variants for RD may share an underlying genetic architecture across population groups for reading disability.

## Introduction

Reading disability (RD) is defined as difficulty in reading despite normal intelligence and adequate instruction. Prevalence ranges from 5% to 17%, and although the relationships between symbols and sounds differ across languages (e.g., alphabetic versus non-alphabetic languages), RD is the most common learning disorder among school-aged children across the world (Ramus, 2003). There is broad consensus that phonological processing deficits play a major role in the etiology of RD (Melby-Lervåg *et al*., 2012; Wagner & Torgesen, 1987). Other contributors include deficits in orthographic processing and processing speed, as well as cognitive and sensory deficits (Caravolas *et al*., 2012; Ramus, 2003; Vidyasagar & Pammer, 2010; Wolf & Bowers, 2000; Wright *et al*., 2000), which are all interrelated at the behavioral level (Wagner *et al*., 1994). Clinically, individuals with double deficits in both phonological processing and processing speed generally have more severe reading problems (Schatschneider *et al*., 2002; Wolf & Bowers, 1999). In addition, there are differences in the underlying neurobiological and genetic factors associated with the above cognitive and behavioral processes (Norton et al., 2015; Pennington et al., 2012). Due to the high frequencies of these shared deficits, Pennington proposed a multiple cognitive deficit model to partially explain the heterogeneous presentations of RD (Pennington, 2006).

Despite the cognitive complexity of RD, researchers have sought to capture component reading skills that contribute to RD in specific groups of individuals using latent profile analysis (LPA), a mixture modeling technique that aims to identify homogenous subgroups of individuals based on similarities in their response patterns to a number of continuous variables (Oberski, 2016). The application of LPA has been successful in describing multiple subtypes of struggling readers with varying strengths and weaknesses across reading and language-related processes, including phonological processing, naming speed, vocabulary, comprehension (listening and reading), and word level skills (fluency, decoding, and accuracy) (Brasseur-Hock *et al*., 2011; Lesaux & Kieffer, 2010; O’Brien *et al*., 2012; Wolff, 2010). Latent profiles of RD that have been identified thus far are dependent on the sample from which the profiles were derived as well as the cognitive variables assessed in the models. Therefore, a majority of the literature focuses on dissecting RD based on component reading-specific skills and processes, but there is limited work that examines RD subtypes based on cross-clinical phenotypes that commonly co-occur with RD including phenotypes associated with attention deficit hyperactivity disorder (ADHD) and language learning disability (LLD).

In addition to co-occurring deficits, there is a growing body of evidence linking RD to other developmental and behavioral disorders. RD is frequently comorbid with both ADHD (McGrath *et al*., 2011; Willcutt, 2012; Willcutt & Pennington, 2000) and LLD (Bishop & Snowling, 2004; McArthur *et al*., 2000; Melby-Lervåg & Lervåg, 2014). The co-occurrence of RD in subjects selected for ADHD is between 25% and 40% (Sexton *et al*., 2012), while the co-occurrence of ADHD in subjects selected for RD is between 15% and 35% (Willcutt *et al*., 2000). The co-occurrence of RD and ADHD is more associated with inattention than with hyperactivity-impulsivity (Plourde *et al*., 2015; Rosenberg *et al*., 2012). Impairment in processing speed is a primary cognitive risk factor that is shared by RD and ADHD (McGrath *et al*., 2011; Shanahan *et al*., 2006; Willcutt *et al*., 2005). In addition to processing speed, phonological processing deficits are a shared risk factor for RD and ADHD comorbidity (Purvis & Tannock, 2000). Language learning disability (LLD) refers to language impairment that is defined by language-based deficits in development of vocabulary and grammar (Bishop *et al*., 2016; Bishop & Snowling, 2004) with difficulty in vocabulary considered a core deficit. The term LLD rather than language impairment is used in this study because it is emerging as a general umbrella term to refer to any language-based disorder (Bishop *et al*., 2016; Bishop & Snowling, 2004). Similar to RD and ADHD, RD and LLD co-occur in 15%-40% of cases (Willcutt & Pennington, 2000). They share many of the same cognitive deficits, including phonological processing and language fluency (Catts *et al*., 2005; Pennington, 2006; Pennington & Bishop, 2009; Wise *et al*., 2007). Given these common comorbidities that could be explained by shared cognitive processes contributing to these clinical disorders, an LPA framework would be appropriate to parse struggling readers with subclinical presentations of ADHD and/or LLD that could contribute to the manifestation of RD in these individuals.

Examination of the genetic effects that contribute to the co-occurrence of RD and ADHD strongly implicate the RD risk gene, DCDC2, located on human chromosome 6p22, suggesting a pleiotropic effect within this region (Scerri *et al*., 2011). A univariate linkage analysis for ADHD and bivariate linkage analysis for ADHD and reading measures in a sample of sibling-pairs selected for co-occurrence identified a significant linkage signal in the region of 6p22(Willcutt *et al*., 2002). This was further supported by genetic analyses variants within *DCDC2* (Meng *et al*., 2011; Meng *et al*., 2005; Powers *et al*., 2013; Powers *et al*., 2016). Quantitative measures of language (Cope et al., 2012), inattention and hyperactivity/impulsivity (Couto et al., 2009; Willcutt et al., 2002) have also been associated with variants of *DCDC2*, as well as a gene-by-gene interaction between *DCDC2* and *KIAA0319*, another candidate risk gene for RD (Mascheretti et al., 2017; Riva et al., 2015). These studies provide compelling evidence that *DCDC2* has pleiotropic effects on reading, language, and attention. We hypothesize that genetic variants from this gene is associated with shared deficits in RD, ADHD and LLD, and could be used to clinically differentiate learners so that they can be provided with the optimal intervention tailored to their individual needs.

Considerable advances have been made in understanding RD from behavioral and genetic perspectives. However, most study designs do not consider the range of heterogeneous clinical presentations of RD and how genetics factors could differentially contribute to each. Furthermore, most research on the genetics of RD has focused on populations of European descent. There are known differences in the genetic architecture of the genome across population groups, raising the possibility that genetic associations previously identified in European samples could be population specific or shared (Li & Keating, 2014). In the present study, we first identify different classes of RD based on a latent profile analysis (LPA) of standard scores from nine reading, language, and attention assessments in a diverse sample of Hispanic American and African American youth. We then investigate whether genetic variants of the READ1 transcriptional modifier for *DCDC2* are associated with specific classes of struggling readers. Due to prior associations and evidence of molecular function, we predict that READ1 will show association with one or more RD classes, providing evidence of a contribution to a shared genetic architecture with potential clinical applications.

## Methods

### Participants

Participants were 1,432 self-identified African American (n = 531) and Hispanic American (n = 955) children, age 8 to 15 years, enrolled between 2010 and 2014. Note that participants could self-identify as both African American and Hispanic, resulting in a small group that overlapped both categories. This study was part of a larger, multi-site US and Canadian collaborative Genes, Reading, and Dyslexia (GRaD) project led by Yale University. The study initially followed a case control design, but recruitment difficulties led to a change in study design to include the full range of reading ability, with heavy oversampling of the low end of reading skill. Participating sites included Albuquerque, NM; Baltimore, MD; Boston, MA; Boulder and Denver, CO; New Haven, CT; San Juan, PR; and Toronto, Canada.

Initial inclusion criteria for RD in the GRaD study were either history of poor reading skills, report of skills falling below expected level for age or grade, and/or provision of special services in reading. Inclusion criteria for controls in the GRaD study were reading skill scores above expectations for grade, performance above the 40th percentile on standardized school, or clinical testing. Exclusion criteria were as follows: age outside the target age range; non-minority ethnic or racial group membership; foster care placement; preterm birth (defined as < 36 weeks gestation); prolonged stay in the NICU after birth (defined as > 5 days); history of diagnosed or suspected significant cognitive delays, significant behavioral problems, or frequent school absences; history of serious emotional/psychiatric disturbances (i.e., major depression, psychotic or pervasive developmental disorder, Autism) or chronic neurologic condition (i.e., seizure disorder, developmental neurological conditions, Tourette or other tic disorders, acquired brain injuries); and documented vision or hearing impairment. This study was approved by the Human Investigation Committee of Yale University and all the review boards of participating data collection sites (University of Colorado-Boulder, University of Denver, Tufts University, University of New Mexico, Kennedy Krieger Institute, and the Hospital for Sick Children-Toronto). Parental consent forms and child assent were collected before participation.

Prior to enrollment, suspected RD and control status was confirmed with individually-administered standardized reading tests. In the present study, RD cases were defined by standard scores below 85 on two or more reading composites (WJ-III, TOWRE, SRI; n = 329). Controls were typical readers who scored better than −1 SD on the two or more reading composites (n = 1103). Among the 329 children with RD, there were 73 African-American and 241 Hispanic-American children (with 15 who did not self-identified). Among 1103 controls, there were 256 African-American and 812 Hispanic-American children (with 35 who did not self-identified).

### Measures

Reading outcome assessments consisted of individually administered standardized measures including Woodcock-Johnson III - Letter-Word Identification and Word Attack (Woodcock *et al*., 2001), Test of Word Reading Efficiency - Sight Word Efficiency and Phonetic Decoding Efficiency (TOWRE) (Torgesen *et al*., 1999), and Standardized Reading Inventory - Word Recognition and Reading Comprehension (SRI) (Newcomer, 1999). Core reading-related skills measures included well-validated instruments tapping phonological awareness assessed by the Comprehensive Test of Phonological Processing – elision and blending (CTOPP) (Wagner *et al*., 1999), naming speed assessed by the Rapid Automatized Naming – letters, numbers, and objects (RAN) (Wolf & Denckla, 2005), language assessed by the Peabody Picture Vocabulary Test (PPVT) (Dunn & Dunn, 2007) and the Wechsler Intelligence Scale for Children (WISC) – vocabulary (Wechsler *et al*., 2004), along with inattention and hyperactivity assessed by the Strengths and Weakness of ADHD symptoms and Normal behavior rating scale (SWAN) (Swanson *et al*., 2006).

#### Woodcock-Johnson Tests of Achievement, Third Edition (WJ-III)

Measures from the WJ-III included the Letter-Word Identification and Word Attack subtests (Woodcock *et al*., 2001). The WJ-III Letter Word Identification subtest is an untimed measure of non-contextual single word reading ability requiring the child to read a list of increasingly complex English words aloud. The Word Attack subtest asks the participant to apply knowledge of English phonology to decode non-words or pseudowords in isolation. The total score for each subtest represents the number of words read correctly, converted to a standard score based upon age norms.

#### Test of Word Reading Efficiency (TOWRE)

The TOWRE is an assessment of the child’s single word reading and single pseudoword decoding isolated (non-contextual) word fluency under timed conditions (Torgesen *et al*., 1999). The child is asked to read as many individual words (Sight Word Efficiency) or non-words (Phonetic Decoding Efficiency) of increasing length and phonetic difficulty as possible in 45 seconds. Scores for Sight Word Efficiency and Phonetic Decoding Efficiency represent the number of correctly read words within the time limit, relative to age norms.

#### Standardized Reading Inventory, Second Edition (SRI)

The SRI is an individually-administered contextual reading test that consists of 10 passages of increasing difficulty, ranging from pre-primer to an eighth-grade level (Newcomer, 1999). Oral reading accuracy is assessed and children are then asked to answer a series of comprehension questions. Scores are obtained for word recognition accuracy and comprehension on each passage; the total score in each skill area is converted to a norm-referenced standardized score.

#### Comprehensive Test of Phonological Processing (CTOPP)

CTOPP is a comprehensive instrument designed to assess phonological awareness, phonological memory, and rapid naming (Wagner *et al*., 1999). In the current study, Elision and Blending Words subtests were used to measure children’s phonological awareness. Elision measures the ability to remove phonological segments from spoken words to form other words. Blending Words measures the ability to synthesize sounds to form words.

#### Rapid Automatized Naming (RAN) and Rapid Alternating Stimulus (RAS) Tests

RAN Letters, RAN Objects, and RAS Letters and Numbers were assessed in the GRaD sample. The Letters and Objects subtests from the RAN assessed speeded lexical retrieval, requiring the child to rapidly name a closed set of 5 letters or objects (repeated randomly in 5 rows of 10 items) as quickly as possible without making mistakes (Wolf & Denkla, 2005). RAS Letters and Numbers is similar in design to RAN, but requires the child to rapidly name stimuli from alternation stimulus categories (alternating letters and numbers). Time to completion is recorded and converted to an age-referenced standard score.

#### Peabody Picture Vocabulary Test, Fourth Edition (PPVT)

The PPVT-4 is an untimed measure of receptive vocabulary knowledge; the participant is required to point to one of four pictures that best indicates the target word presented (Dunn & Dunn, 2007). Total raw scores (correct responses) are converted to age-normed standardized scores.

#### Wechsler Intelligence Scale for Children, Fourth Edition (WISC-IV)

The WISC-IV Vocabulary measures children’s verbal fluency, concept information, word knowledge, and word usage (Wechsler *et al*., 2004). Children were asked to give oral definitions of words. Scoring was 2-1-0, according to the quality of the responses.

#### Strengths and Weakness of ADHD symptoms and Normal behavior rating scale (SWAN)

The SWAN has 18 ADHD items based on DSM-5 criteria for inattention and hyperactivity/impulsivity symptoms (Swanson *et al*., 2006). The first 9 items assess inattention symptoms, and the remaining assess hyperactivity/impulsivity symptoms. The Seven-Likert scale from “far below” to “far above” was used to rate children’sbehavior by the parent.

### DNA collection and genotyping

Saliva was collected using Oragene-DNA kits (DNA Genotek Inc.) and DNA extracted with prepIT-L2P (DNA Genotek Inc.) using manufacturer protocols. Individuals were removed from analysis if there were discrepancies between self-reported sex and inferred sex based on X-chromosome heterozygosity (Thornton *et al*., 2012). READ1 genotyping was conducted using PCR amplification and Sanger sequencing. Sanger sequencing was performed at the Yale W.M. Keck DNA Sequencing Facility using standard protocols. Primer sequences and amplification protocol are described in detail elsewhere (Powers *et al*., 2013). READ1 alleles were called from chromatograms using a custom program written in C++ (Dr. Yong Kong, available upon request). If the calling program identified errors, chromatograms were manually examined and deconvoluted for allele calling. The genotyping call rate for READ1 alleles was 0.987.

Genotyping for the 2,445 bp microdeletion on 6p22, which encompasses the READ1 allele within its breakpoints, was conducted using allele specific PCR and agarose-gel electrophoresis. Primer sequences and amplification protocol for microdeletion genotyping are described in detail elsewhere (Powers *et al*., 2013). In brief, the three-primer reaction generates a ∼600 bp amplicon when the microdeletion is not present, and a ∼200 bp amplicon when the microdeletion is present. The PCR products were resolved by electrophoresis through a 1% agarose gel with ethidium bromide (0.2 µg/mL) in 1X TBE buffer run at 125V for 60 minutes. Gels were visualized using a UV transilluminator (Molecular Imager Gel Doc XR, Bio-Rad Laboratories) and genotypes were manually called from the gels. The microdeletion genotyping call rate was 0.972.

### Functional Groups of READ1 Alleles

Three functional groups of READ1 alleles were investigated (Table 1). RU1-1 alleles have only one copy of Repeat Unit 1 (RU1-1: alleles 2, 3, 9, 12, 25, 27). RU1 sequence was previously used to capture ETV6, a dimerizing transcriptional repressor that binds to a consensus binding site, GGAAG, which is present in the RU1 site: GAGAGGAAGGAAA. The RU1-1 group of alleles was associated with a moderate protective effect on reading performance in the Avon Longitudinal Study of Parents and Children (ALSPaC), a longitudinal cohort of European descent (Powers et al., 2016). RU1 also contains a consensus binding site (GAGAGGAAGGAaagg) for many C2H2 domains that define classical zinc finger transcription factors.

**Table 1.** Functional grouping of READ1 alleles and previously reported associations

| Functional Group | Definition | Alleles | Phenotype |
| --- | --- | --- | --- |
| RU1-1 | One copy of Repeat Unit 1 | 2, 3, 8, 12, 25, 27 | Protective effect for RD in ALSPaC (Powers *et al.*, 2016) |
| RU2-Long | $\geq 8$ copies of Repeat Unit 2 | 5, 6, 13, 14, 19, 20, 22, 23 | Poor reading performance in ALSPaC (Powers *et al.*, 2016) |
| RU2-Short | $\leq 6$ copies of Repeat Unit 2 | 4, 10, 16, 21 | Poor reading comprehension in GRaD (Li *et al.*, 2018) |

RU2-Long alleles have two copies of RU1 and greater than seven copies of Repeat Unit 2 (RU2: alleles 5, 6, 13, 14, 19, 20, 22, 23; Table 1). RU2 (GGAA) contains consensus-binding sites for ETV6. In previous studies, allele 5 (RU2-long) was associated with increased risk for RD, whereas allele 6 (RU2-long) was associated with increased risk for LLD (Powers *et al*., 2016).

RU2-Short is characterized by alleles that have fewer than six copies of RU2 (alleles 4, 10, 16, 21). The most frequent alleles in this category are alleles 4 and 10 which did not show association with either a positive or negative impact on reading performance in our previous studies with the ALSPAC (Powers *et al*., 2013; Powers *et al*., 2016). The sequence of READ1 alleles is presented in Table 2.

**Table 2.** Sequence of READ1 alleles

| RU1-1 |  |  |  |  |  |  |  |  |
| --- | --- | --- | --- | --- | --- | --- | --- | --- |
| Allele | Repeat unit 1 | Repeat unit 2 | SNP1 | Repeat unit 3 | Const. Region | Repeat unit 4 | Repeat Unit 5 | Length (BPs) |
| 2 | (GAGAGGAAGGAAA) 1 | (GGAA) 9 | (GAAA) 0 | (GGAA) 0 | GGAAAGAATGAA | (GGAA) 4 | (GGGA) 2 | 85 |
| 3 | (GAGAGGAAGGAAA) 1 | (GGAA) 6 | (GAAA) 1 | (GGAA) 2 | GGAAAGAATGAA | (GGAA) 4 | (GGGA) 2 | 85 |
| 9 | (GAGAGGAAGGAAA) 1 | (GGAA) 7 | (GAAA) 1 | (GGAA) 2 | GGAAAGAATGAA | (GGAA) 4 | (GGGA) 2 | 89 |
| 12 | (GAGAGGAAGGAAA) 1 | (GGAA) 8 | (GAAA) 1 | (GGAA) 2 | GGAAAGAATGAA | (GGAA) 3 | (GGGA) 2 | 89 |
| 25 | (GAGAGGAAGGAAA) 1 | (GGAA) 8 | (GAAA) 1 | (GGAA) 2 | GGAAAGAATGAA | (GGAA) 4 | (GGGA) 2 | 93 |
| 27 | (GAGAGGAAGGAAA) 1 | (GGAA) 5 | (GAAA) 1 | (GGAA) 2 | GGAAAGAATGAA | (GGAA) 4 | (GGGA) 2 | 81 |
| RU2-Long |  |  |  |  |  |  |  |  |
| Allele | Repeat unit 1 | Repeat unit 2 | SNP1 | Repeat unit 3 | Const. Region | Repeat unit 4 | Repeat Unit 5 | Length (BPs) |
| 5 | (GAGAGGAAGGAAA) 2 | (GGAA) 8 | (GAAA) 1 | (GGAA) 2 | GGAAAGAATGAA | (GGAA) 4 | (GGGA) 2 | 106 |
| 6 | (GAGAGGAAGGAAA) 2 | (GGAA) 8 | (GAAA) 1 | (GGAA) 2 | GGAAAGAATGAA | (GGAA) 3 | (GGGA) 2 | 102 |
| 13 | (GAGAGGAAGGAAA) 2 | (GGAA) 9 | (GAAA) 1 | (GGAA) 2 | GGAAAGAATGAA | (GGAA) 3 | (GGGA) 2 | 106 |
| 14 | (GAGAGGAAGGAAA) 2 | (GGAA) 9 | (GAAA) 1 | (GGAA) 2 | GGAAAGAATGAA | (GGAA) 4 | (GGGA) 2 | 110 |
| 19 | (GAGAGGAAGGAAA) 2 | (GGAA) 9 | (GAAA) 0 | (GGAA) 0 | GGAAAGAATGAA | (GGAA) 4 | (GGGA) 2 | 98 |
| 20 | (GAGAGGAAGGAAA) 2 | (GGAA) 10 | (GAAA) 1 | (GGAA) 2 | GGAAAGAATGAA | (GGAA) 4 | (GGGA) 2 | 114 |
| 22 | (GAGAGGAAGGAAA) 2 | (GGAA) 10 | (GAAA) 0 | (GGAA) 0 | GGAAAGAATGAA | (GGAA) 4 | (GGGA) 2 | 102 |
| 23 | (GAGAGGAAGGAAA) 2 | (GGAA) 11 | (GAAA) 0 | (GGAA) 0 | GGAAAGAATGAA | (GGAA) 4 | (GGGA) 2 | 106 |
| RU2-Short |  |  |  |  |  |  |  |  |
| Allele | Repeat unit 1 | Repeat unit 2 | SNP1 | Repeat unit 3 | Const. Region | Repeat unit 4 | Repeat Unit 5 | Length (BPs) |
| 4 | (GAGAGGAAGGAAA) 2 | (GGAA) 6 | (GAAA) 1 | (GGAA) 2 | GGAAAGAATGAA | (GGAA) 4 | (GGGA) 2 | 98 |
| 10 | (GAGAGGAAGGAAA) 2 | (GGAA) 4 | (GAAA) 1 | (GGAA) 2 | GGAAAGAATGAA | (GGAA) 4 | (GGGA) 2 | 90 |
| 16 | (GAGAGGAAGGAAA) 2 | (GGAA) 5 | (GAAA) 1 | (GGAA) 2 | GGAAAGAATGAA | (GGAA) 4 | (GGGA) 2 | 94 |
| 21 | (GAGAGGAAGGAAA) 2 | (GGAA) 6 | (GAAA) 1 | (GGAA) 2 | GGAAAGAATGAA | (GGAA) 3 | (GGGA) 2 | 94 |

Note. The number adjacent to the parentheses indicates the number of times this sequence is repeated.

#### Population Stratification Correction

A principal components analysis (PCA) on genome-wide genotyped SNPs (Illumina Infinium Omni2.5) was conducted to model continuous axes of genetic variation within the GRaD sample using EIGENSTRAT and is a common method used to correct for differences in genetic substructure across ethnic groups (Cook & Morris, 2016; Price *et al*., 2006). Prior to analysis, SNP level quality control included the removal of SNPs with missingness greater than 5%, Hardy-Weinberg equilibrium p<0.0001, were not autosomal and had a minor allele frequency less than 0.05. Sample level quality control included the removal of samples with percentage of missing genotypes greater than 3% (n = 39) and samples with discrepancies between reported and inferred sex based on X chromosome heterozygosity (n = 52). The first 10 principal components generated were used to correct for genomic inflation due to allele frequency differences across different ancestries. Analyses of genome-wide SNP data in the GRaD study found that the first 10 principal components were effective in reducing the effects of population stratification (genomic inflation factor < 1.05) in the phenotypes evaluated in the present analysis (Truong *et al*., 2017). Furthermore, self-reported ethnicity is consistent with the PCA analysis based on the first two principal components (Supplemental Figure 1).

### Reading disability identification criteria

In the present study, subjects with RD (n = 329) were identified as children whose standard scores were below 85 (i.e., < 1 SD) on two or more reading composites (WJ-III, TOWRE, SRI). Controls (n = 1103) were typical readers who scored better than a standard score of 85 on the two or more reading composites. These definitions were used to inform the latent profile analysis.

### Statistical Analysis

#### Latent Profile Analysis

Latent profile analysis (LPA) using M-plus (Muthén & Muthén, 2012) was conducted to identify subgroups of typical and poor readers based on standard scores from nine reading, language and attention indicators, which were all continuous variables. In the past, ‘cut-offs’ were typically used to identify a person with RD. An individual who scores below certain cut-offs (i.e. the 25th percentile on decoding and/or word reading fluency and/or phonological tasks) is considered to have a RD. However, it is unclear whether the people around the 25th boundary would be classified with RD or not. The advantage of LPA is that it can identify naturally occurring clusters of poor readers without depending on poor reading performance cut-offs (Volk et al., 2006). In the present analysis, the nine indicators were PPVT, CTOPP elision, CTOPP blending, WISC vocabulary, RAN objects, RAN letter, RAN number, SWAN inattention, and SWAN hyperactivity.

Model fit was evaluated by several fit indices: Akaike Information Criterion (AIC), Bayesian Information Criterion (BIC), Entropy, and Lo-Mendell-Rubin Adjusted Likelihood Ratio Test (LMR-LRT). These four indices are the most widely accepted indices when reporting LPA results (Nylund *et al*., 2007). The model with the smallest values on AIC and BIC is considered to have the best fit. Entropy ranging from 0 to 1 is the degree of classification accuracy. Higher entropy values reflect better classification(Tein *et al*., 2013). LMR is used to compare models with different numbers of classes. A non-significant value suggests that the model with one fewer class is a more concise representation of the data.

#### Logistic Regression with RD classes as Outcome

The primary multivariate evaluation of the latent classes was conducted via three logistic regressions using raw scores. In each regression, one individual RD latent class (e.g., either 4, 5, or 6) was contrasted against three non-RD latent classes (e.g., 1, 2 and 3). The three RD classifications were regressed onto a suite of control variables and subsequently onto the READ1 deletion and allele groups as predictors. The first step of control variables consisted of the ten principal components to control for population stratification. The second step of control variables consisted of the following: child’s age in months; whether Spanish was the primary language spoken in the home at the time of the child’s birth, whether parents reported a diagnosis of ADHD for their child; the highest level of education reported for the primary caretaker; and, whether the family had used any in a list of government social assistance programs (e.g., food stamps, Medicaid, federal WIC, etc.). These variables have been shown to be highly associated with reading ability (Bradley & Corwyn, 2002; Hart *et al*., 2013; Snowling & Hulme, 2008). The third step of genetic variables consisted of the following: presence of READ1 RU1-1, READ1 microdeletion, RU2-Short, and RU2-Long allele groups. Subjects with mixed READ1 allele groups were coded as “present” in both categories (i.e. categorized as “present” under the RU1-1 and RU2-Long groups, respectively), and included in the logistic regression.

The goal of this analysis was to ascertain whether the genetic variants predicted each of the three RD latent classes, controlling for common phenotypic and environmental factors previously found to be associated with RD. Across the three logistic regression models, we used a base alpha of .05, with a correction to control the false-discovery rate (Benjamini & Hochberg, 1995) and prevent overidentification of individual predictors among covariates and genetic factors (Benjamini & Hochberg, 1995).

## Results

### Characterization of Reading Outcomes for the RD and Control Groups

Table 3 shows the performance (age standardized scores) of reading outcomes and reading-related measures for the RD and control groups. This illustrates that individuals within the RD group had poor performance across several different reading measures, while individuals in the control group showed proficient performance. In addition, the definition of RD groups by replication across two composite measures resulted in a homogeneous and well-defined group, as evidenced by the smaller standard deviation.

**Table 3.** Means and standard deviations for RD and control groups on reading outcome and reading-related measures

|  |  | Mean<br>(Standard Deviation) |  |  |  |  |  |
| --- | --- | --- | --- | --- | --- | --- | --- |
|  |  | WJ<br>LWI | WJ<br>WA | TOWRE<br>SW | TOWRE<br>PD | SRI<br>WR | SRI<br>RC |
| RD |  | 78.99<br>(11.53) | 81.80<br>(9.35) | 78.07<br>(8.84) | 75.27<br>(7.97) | 3.12<br>(1.67) | 4.40<br>(2.30) |
| Control |  | 100.92<br>(10.46) | 98.77<br>(8.98) | 100.29<br>(10.60) | 99.60<br>(12.57) | 8.41<br>(3.71) | 8.55<br>(3.65) |

|  |  | Mean<br>(Standard Deviation) |  |  |  |  |  |  |  |  |
| --- | --- | --- | --- | --- | --- | --- | --- | --- | --- | --- |
|  |  | PPVT | WISC<br>Vocabulary | CTOPP<br>Elision | CTOPP<br>Blending | RAN<br>Objects | RAN<br>Letters | RAS<br>Letters&Numbers | SWAN<br>Inattention | SWAN<br>Hyperactivity |
| RD |  | 85.15<br>(12.86) | 6.63<br>(2.48) | 5.72<br>(2.32) | 7.58<br>(2.39) | 88.67<br>(14.28) | 90.79<br>(12.29) | 89.74<br>(12.73) | 3.08<br>(10.17) | -1.23<br>(10.66) |
| Control |  | 97.44<br>(15.16) | 9.47<br>(2.82) | 9.23<br>(2.65) | 9.70<br>(2.49) | 99.48<br>(15.31) | 103.77<br>(13.08) | 103.90<br>(12.67) | -3.56<br>(11.27) | -5.61<br>(11.38) |

*Note.* WJ LWI = Woodcock-Johnson Tests of Achievement - Letter Word Identification, WJ WA = Woodcock-Johnson Tests of Achievement - Word Attack; TOWRE SW = Test of Word Reading Efficiency - Sight Word; TOWRE PD = Test of Word Reading Efficiency – Phonemic Decoding; SRI WR = Standardized Reading Inventory – Word Recognition; SRI RC = Standardized Reading Inventory – Reading Comprehension; PPVT = Peabody Picture Vocabulary Test; WISC = Wechsler Intelligence Scale for Children; CTOPP = Comprehensive Test of Phonological Processing; RAN = Rapid Automatized Naming; RAS = Rapid Alternating Stimulus; SWAN = Strengths and Weakness of ADHD symptoms and Normal behavior rating scale. Age-standardized normative expected values for WJ-III, TOWRE, PPVT, WISC, and RAN have a mean of 100, with a standard deviation of 15; Age-standardized normative values for the SRI and CTOPP have a mean of 10, with a standard deviation of 3.

### Latent Profile Analysis

Table 4 shows the results of models for each class solution. Models with two through four latent classes were compared to select a best fit. All four indices (AIC, BIC, Entropy, and Lo-Mendell-Rubin) reflecting model fit indicated that the 3-class model was the optimal model that provides the best representation of the reading-related performance in the GRaD sample given the input variables.

**Table 4.** Indicators of fit for models with two though four latent classes

| Model | AIC | BIC | Entropy | Lo-Mendell-Rubin |
| --- | --- | --- | --- | --- |
| 2-class | 83893.096 | 84087.579 | 0.772 | 2113.065<br>($p < 0.001$) |
| 3-class | 83093.041 | 83387.394 | 0.801 | 832.02<br>($p = 0.010$) |
| 4-class | 82706.874 | 83101.096 | 0.776 | 421.113<br>($p = 0.5337$) |

AIC: Akaike Information Criterion
BIC: Bayesian Information Criterion
Lo-Mendell-Rubin: Adjusted Likelihood Ratio Test

The initial RD/non-RD assignments were also included as a known-class variant (Lubke & Muthén, 2005) to estimate the 3-class model with multiple groups (Lubke & Muthén, 2005). As a result, six latent classes were identified in total: three typical reading latent classes (Classes 1-3), and three RD latent classes (Classes 4-6). Figure 1 utilizes a violin plot to depict the relative profile of skills across the input measures, within each class. Class 1 (*n* = 314) had exceptional good reading and performed well on all cognitive, linguistic, and attention assessments. Class 2 (*n* = 367) had average reading but was relatively poor on naming speed. Class 3 (*n* = 414) had average reading but was relatively low on vocabulary. Within the RD latent classes, Class 4 (*n* = 142) and Class 5 (*n* = 118) had relative strength in vocabulary and phonology, but Class 4 was the most deficient in naming speed and phonological blending, matching the profile of co-occurring RD+ADHD. Class 5 shared the reading impairment of Class 4 and Class 6. This class appeared to have only relative weakness in phonological awareness, with vocabulary, rapid naming, phonological awareness and attention *relatively* intact compared to the other two groups. Class 5 thus resembles RD with specific phonological impairment. Class 6 (*n* = 63) had the lowest vocabulary and phonological awareness, but relative strength in rapid naming, and matched the profile of co-occurring RD+LLD. Figure 1 displays the performance of different classes across all nine reading, language, and attention measures. Supplemental Table 1 in the *Supporting Information* presents the estimated means and standard errors for each class.

**Figure 1:**
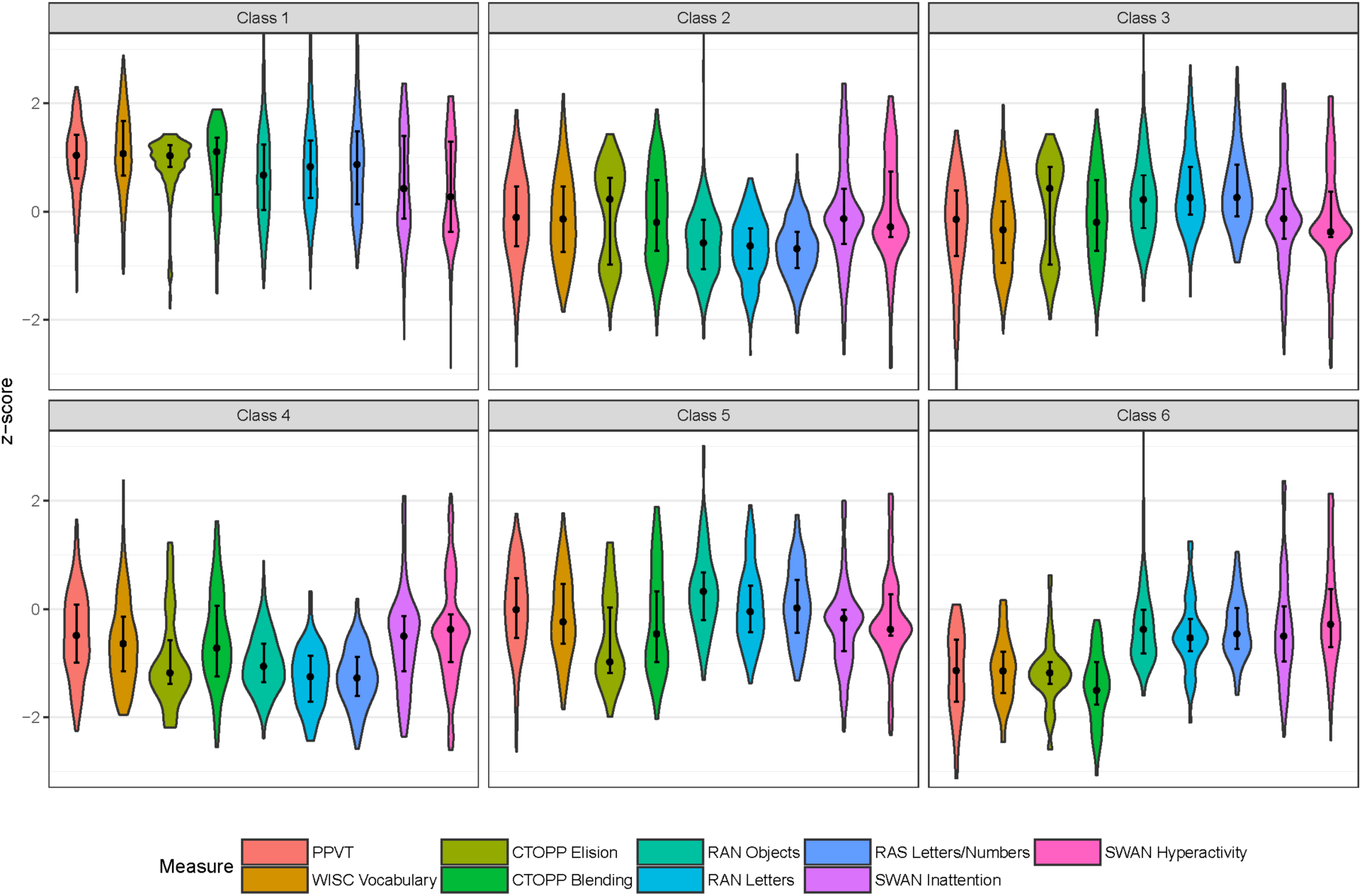
Violin plot depicting performance across reading-related measures for each of the six latent profiles. Dots represent median values, while straight lines above and below the median values represent the interquartile range.

### Associations between RD latent classes and READ1 allele groups

The primary association analysis consisted of three logistic regressions, each contrasting one RD latent class against the three non-RD classes (e.g., Class 1, 2 and 3 from the LPA), while controlling for population stratification and covariates. The first ten principal components were entered as a block in the first step and accounted for a significant and moderate percent of the variance in each of the three RD classes (Class 4, Nagelkerke *R^2^* = .045; Class 5, *R^2^* = .043; Class 6, *R^2^* = .094).

Table 5 summarizes the results of the three logistic regressions. SES, child’s age, Spanish spoken at home, previous ADHD diagnosis, and caretaker’s education level were covariates because these variables are believed to be influential factors associated with reading ability. When LPA Class 4 (RD plus shared ADHD features: relatively poor on phonological awareness but not language, and with profound naming speed and attention problems) was the outcome, child’s age and a diagnosis of ADHD were the only two significant covariates. Controlling for these and all other covariates, RU1-1 was a significant and strong predictor of membership in this latent class (beta = .68 [95% BCa .12 to 1.22], SE = .25, Odds Ratio = 1.98). The presence of RU1-1 contributed almost a doubling of the odds of being in this latent class characterized by both RD and ADHD behavioral features.

**Table 5.** Prediction of LPA Class 4, 5, and 6 from covariates, READ1 deletion and READ1 functional groups of alleles

|  | Predictor | Odds Ratio (95% Bootstrap CI) |  |  |
| --- | --- | --- | --- | --- |
|  |  | Class 4 vs. (1, 2, 3) | Class 5 vs. (1, 2, 3) | Class 6 vs. (1, 2, 3) |
| Covariates | SES | 1.45<br>(0.86, 2.34) | 1.57<br>(0.85, 3.02) | **2.68**<br>**(1.27, 6.17)** |
|  | Child's Age | **0.98**<br>**(0.97, 0.99)** | **1.03**<br>**(1.02, 1.04)** | **1.03**<br>**(1.01, 1.04)** |
|  | Spanish Spoken at Home | 0.95<br>(0.49, 1.69) | 0.96<br>(0.41, 1.81) | **2.81**<br>**(1.03, 8.75)** |
|  | Prev. ADHD Diagnosis | **2.81**<br>**(1.81, 5.16)** | 1.64<br>(0.81, 2.86) | 1.41<br>(0.48, 3.56) |
|  | Caretaker Education | 0.95<br>(0.85, 1.08) | 1.03<br>(0.94, 1.14) | 1.02<br>(0.90, 1.17) |
| Genetic Variants | READ1 RU1-1 | **1.98**<br>**(1.21, 3.25)** | 1.64<br>(0.86, 3.31) | 0.98<br>(0.19, 2.07) |
|  | READ1 RU2-Short | 1.32<br>(0.71, 2.04) | 1.14<br>(0.59, 2.02) | 1.58<br>(0.58, 3.52) |
|  | READ1 RU2-Long | 0.79<br>(0.39, 1.71) | 1.03<br>(0.42, 2.14) | 1.90<br>(0.67, 5.24) |
|  | READ1 Microdeletion | 0.95<br>(0.24, 2.01) | **2.27**<br>**(1.08, 4.36)** | 2.17<br>(0.40, 6.51) |

*Note.* Bold values indicate that estimates are significant at .05 level under bootstrap distribution; 10 principal components (population stratification correction) were entered in the first step prior to covariates and genetic variants but are not reported here.

When LPA Class 5 (RD behavioral features: specific relative weakness in phonological awareness) was the outcome, child’s age was the only significant covariate. Controlling for age and all covariates, the READ1 microdeletion significantly predicted class membership (beta = .82 [95% BCa .06 to 1.50], SE = .33, Odds Ratio = 2.27), doubling the odds of being in this RD latent class.

When LPA Class 6 (RD with LLD features: global language and phonological awareness problems, but intact attention and naming speed) was the outcome, low SES, increasing child’s age, and Spanish home language were associated with a higher probability of being in this latent class. No READ1 markers significantly predicted membership in this RD class.

Two follow-up sensitivity analyses were conducted to determine the robustness of the initial results. In the first, each RD LPA class was compared to all other classes (e.g., 5 vs. 1, 2, 3, 4, & 6 together). Across all three logistic regressions, the same pattern of results was obtained, though slightly weaker (e.g., the odds-ratio representing the effect of RU1-1 on Class 4 was 1.86, compared to 1.98 when Class 4 was compared to only the RD latent classes as reported in Table 5 above). In the second sensitivity analysis, each of the RD latent classes (4, 5, and 6) were compared with Class 1 alone, the only non-RD class that did not have *relative* weakness on any cognitive or linguistic skill. Across all three logistic regressions, the same pattern of results was obtained as the main analyses reported above, though again with reduced effect.

## Discussion

Our classification model inputs were theoretically driven by a model that included core RD characteristics plus commonly co-occurring language and attention deficits. The predictors included in the model were as follows: phonological awareness, processing speed, vocabulary, and attention. Extensive evidence has supported the crucial role of phonological awareness in understanding RD, and phonological core deficits are a strong predictor of RD (Melby-Lervåg *et al*., 2012; Wagner & Torgesen, 1987). Processing speed is another powerful predictor of RD (Schatschneider *et al*., 2002; Wolf & Bowers, 1999). The double-deficit theory suggests that deficits in both phonological awareness and processing speed could result in the most severe difficulties with reading. Vocabulary has been shown to be one of the most important predictors of word reading and reading comprehension, and is a main determining factor for presence or absence of LLD (Bishop *et al*., 2004; Catts *et al*., 2005). Inattention was included because, along with processing speed, it is a replicated behavioral marker of co-occurring RD+ADHD (Shanahan *et al*., 2006,;Willcutt *et al*., 2005). Similarly, phonological awareness has also been shown to be a cognitive indicator of co-occurring RD+LLD (Catts *et al*., 2005; Pennington & Bishop, 2009; Wise *et al*., 2007). Therefore, the inclusion of these factors was theory-driven and captured overall representations of RD, LLD, ADHD, and their co-occurring states.

Latent profile analysis provided a method for reclassifying a heterogeneous population of subjects into homogeneous classes. The interaction of known RD assignment and the nine predictors in the LPA model produced a better understanding of RD subtypes and models of co-occurrence. Each latent class represented a unique pattern of readers. In the six latent classes generated by the LPA model, the first three fell into the control category, which consisted of typical readers. Class 1 readers had the best performance among typical reading classes, excelling across all reading skills. They also did not manifest problems in inattention and hyperactivity. Class 2 and 3 readers had similar average reading skills with the exception of Group 2, which were poorer on RAN, and Group 3 which had lower vocabulary. The performance of these first three typical reading classes demonstrated different types of typical readers in relation to language skill and processing speed. However, we were more interested in the patterns among RD groups.

LPA identified three distinctive classes of RD subjects. Class 4 readers had the most serious inattention and hyperactivity problems, and were therefore considered to represent a co-occurring RD+ADHD. Consistent with results from previous studies, our findings also showed that Class 4 readers had severe deficits in processing speed. These deficits are considered to be a risk factor among individuals who have both RD and ADHD (McGrath *et al*., 2011; Shanahan *et al*., 2006; Willcutt *et al*., 2005). Class 5 readers had relative strength in reading skills among those with RD, but also displayed poor phonological awareness and mild attention issues. As such, this group may represent primary RD in the absence of profound co-occurring language or cognitive deficits. Class 6 readers had the lowest scores across two measures of vocabulary, both receptive and expressive, which suggested co-occurring RD+LLD, showing additional difficulties on phonological awareness tasks. This supports existing studies that have found shared phonological processing deficits in RD and in LLD (Catts *et al*., 2005; Pennington & Bishop, 2009). In line with previous research findings, inattention was found to be more pronounced than hyperactivity-impulsivity in all three RD classes (Plourde et al., 2015; Rosenberg et al., 2012).

The current diagnostic criteria for identifying RD are arbitrary and limited to behavioral domains (Pennington, 2006) generally defined by cut-off criteria. Typically, a child who scores below the 20th to 30th percentile on decoding and/or word reading fluency and/or phonological tasks is considered to have RD, leaving borderline cases either unclassified or forced into a diagnostic category (Branum-Martin *et al*., 2013; Fletcher *et al*., 2006). This limitation to the behavioral cut-off criteria and method for identifying RD derives directly from reliance on linear methods and single constructs (Branum-Martin *et al*., 2013). LPA, in contrast, is optimized to create homogeneous groups based on a broader range of relevant diagnostic indicators (Muthén, 2002). While children are still assigned probabilistically to one group or other, homogeneous groupings reflective of the specific RD subtype may result in more reliable classification.

The association of LPA classes with genetic variants could provide useful information for understanding potential etiological mechanisms underlying RD and co-occurrence with ADHD and LI. RU1-1 of READ1 in *DCDC2* was significantly associated with impairments in processing speed and attention co-occurring with RD (class 4) in the GRaD sample, even after controlling for the significant effects of age and ADHD diagnosis. The results support prior evidence for a pleiotropic role of *DCDC2* in reading and attention performance (Couto *et al*., 2009; Mascheretti *et al*., 2017; Riva *et al*., 2015). To date, there has only been one study that has examined the effects of different READ1 alleles on attention measures. In a study conducted by Riva and colleagues (2015), marginally significant effects of READ1 allele 4 (under the classification of RU2-Short in the present study) were observed with a measure of inattention. However, it is important to note that co-occurrence with poor reading performance was not considered in their analyses, and could potentially explain the differences in observed associations with READ1 alleles.

The reported effects of RU1-1 are more complex in the context of reading performance across different populations. In Western Europeans, Powers et al. (2016) observed a protective effect of RU1-1 when tested for association against severe RD status (Powers *et al*., 2016). In an Italian sample, Trezzi et al. (2017) found no effect of RU1-1 on different reading measures collected in Italian. In the present analysis, the observed effects of RU1-1 were deleterious in Hispanic and African American subjects, though specifically for co-occurring RD with poor processing speed and attention. Taken together, these studies highlight the importance of studying the genetic architecture of RD in different population groups. Hispanic Americans and African Americans have different genetic backgrounds from Europeans, although they both share European background due to recent admixture. Western Europeans and Italians also have slightly different genetic backgrounds but have large differences in orthographic transparency between their languages (English having one of the most opaque and Italian having one of the most transparent). Because of these differences in genetic background and orthographic transparency, there may be variations in the regulatory effects of genetic variants like READ1, due to interactions with ancestry specific variants in *DCDC2*, and other previously unidentified genes and/or regulatory elements that play a part in generating these phenotypes (Meng *et al*., 2011; Powers *et al*., 2016). Therefore, it is possible for RU1-1 alleles to have protective effects on a phenotype in one population, but deleterious effects in another. While we have not yet identified the other variants that help confer these differences, identifying these opposing effects of RU1-1 alleles will aid in the discovery of additional factors contributing to these complex traits. The present findings, together with those from Powers et al. (2016), and Trezzi et al. (2017), highlight that differences in effect of the RU1-1 group of READ1 alleles exist and warrant further investigation to understand why.

The class 5 RD subtype with impairments in reading skills and phonological awareness was associated with the microdeletion of READ1 in *DCDC2*. In general, genetic variants identified in *DCDC2* have previously shown strong and specific associations with word reading and reading disability. However, for the *DCDC2* microdeletion, associations with reading disability are inconclusive. Previous reports from independent German and Italian samples have shown a relationship between the microdeletion and word reading and RD (Marino *et al*., 2011; Wilcke *et al*., 2009), but in other independent studies from Germany, the United Kingdom, and Hong Kong, no associations were observed (Harold *et al*., 2006; Powers *et al*., 2016; Scerri *et al*., 2017). Discrepancies across studies could be attributed to potential differences in the genetic architecture contributing to reading disability across population groups, but larger, more diverse samples, especially those of Hispanic and African ancestry, must be examined to further clarify associations observed with the *DCDC2* microdeletion.

The RD subtype that closely matched the behavioral presentation of co-occurring RD+LLD (class 6) was not associated with READ1 deletion or any READ1 variants. However, in the present study, the logistic regression analysis showed a significant and strong covariate effect of participation in a government assistant program (SES) and exposure to a dual language home environment (Spanish spoken at home) in the co-occurring RD + LLD group. Research has shown that children raised in a low SES environment show disparities in their language development relative to their higher SES peers, with differences in language processing and vocabulary skills observed as early as 18 months of age and persisting into school age (Fernald *et al*., 2013; Perkins *et al*., 2013). There is also evidence that children living in a Spanish-English dual language environment score 1 to 2 SD lower relative to their monolingual peers on measures of expressive and receptive vocabulary and verbal short-term memory prior to enrollment in pre-school (Hammer *et al*., 2008; Mancilla-Martinez & Lesaux, 2011; Paez *et al*., 2007). Although these children make substantial gains in their language development with adequate instruction, they still do not perform as well as their monolingual peers in vocabulary and verbal short-term memory by the end of preschool, with the effects persisting to at least 11 years old (Hammer *et al*., 2008; Mancilla-Martinez & Lesaux, 2011; Paez *et al*., 2007). Failure to attain age-appropriate skills in vocabulary and verbal short-term memory could have deleterious effects on reading comprehension and ability into the school years, which could look like co-occurring RD + LLD (Mancilla-Martinez & Lesaux, 2011). In the present study, it is possible that the co-occurring RD + LLD group is capturing the individuals who have lower SES and live in a dual language environment, and reflect a non-genetic etiology for RD + LLD in our sample.

The LPA approach to identify more homogenous classes of RD and subsequent genetic analysis offers promise in the development of pre-diagnostic screeners for RD that could identify children at risk for specific reading difficulties early before reading delays manifest. Waiting for the presence of low reading performance is problematic and can make intervention efforts more difficult given the severity of the impairments. Research has shown that almost 75% of children can achieve reading skill at an appropriate grade level if they receive intervention within the first three years (Lovett *et al*., 2017). Early intervention before the onset of significant reading delays can have a more positive lasting effect on student outcomes and academic achievement compared to late intervention (Scammacca *et al*., 2007; Wanzek & Vaughn, 2007). Overall, the results suggest the need for granular diagnostic assessment (examining both genetic and non-genetic risk factors) that could inform differentiated intervention.

The current study offers an important initial step in classifying RD into more homogenous subgroups for genetic analysis that could contribute to informative pre-diagnostic screeners for RD. There are several limitations that should be noted. When we assessed language performance, we only relied on two vocabulary measures. Other measures such as morphosyntactic or nonword/sentence repetition tasks should be included to better tap the language factors. The lack of comprehensive measures may explain why genetics associations were not observed in the RD+LLD group as it may not entirely represent the nature of that group. While we were well-powered to conduct a candidate variant approach focusing on genetic variants of known molecular function, we were not well powered to conduct a larger scale genome-wide analysis due to the small sample size within RD classes (n = 63-142). The overall study would benefit from a larger sample and is important for future study to broaden the analysis using a genome-wide approach to identify novel variants contributing to specific subtypes of RD. In the present study, we included children across a broad age range (8-15 years) to better understand RD and its comorbidity across childhood. However, denser sampling and longitudinal follow-up studies are needed to provide more accurate and specific periods of development and better define developmental trajectories.

## Conclusions

To conclude, the findings presented from this empirical study is the first to integrate behavioral and genetic results to characterize subtypes of RD plus co-occurring groups focusing on language and attention. The results bridge the gap between behavioral phenotypes and genetic risk variants, ultimately leading to a more comprehensive understanding of impairments in reading performance. Our study presented empirical results on the understanding of different subtypes of RD and models of co-occurring impairments from cognitive and genetic perspectives in a novel sample of African Americans and Hispanic Americans. However, future studies are needed to explore RD and co-occurring impairments in other populations and orthographies to evaluate the presence of shared and/or independent genetic architecture across populations.

## Acknowledgements

We are indebted to the members of the GRaD Study: the participants and their families, the teams who performed recruiting and testing, the project managers at each site. We would also like to thank the Yale W.M. Keck Biotechnology Resource Laboratory’s DNA Sequencing Resource for Sanger sequencing services.

---

**Supplemental Figure 1:**
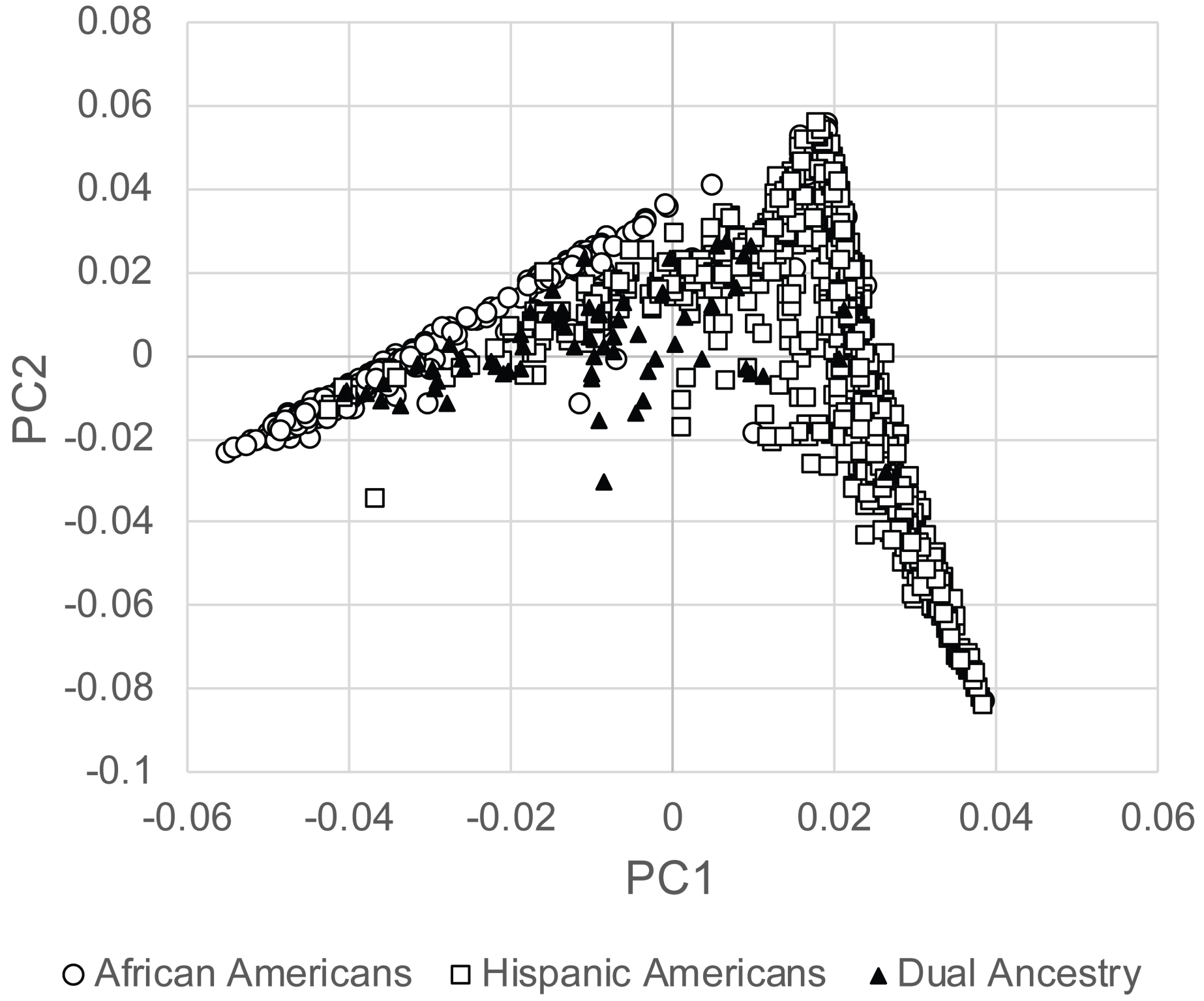
Scatterplot of the first two principal components (PC1 and PC2) against self-reported ancestry (African American, Hispanic American, or Dual Hispanic and African American).

**Supplemental Table 1.**
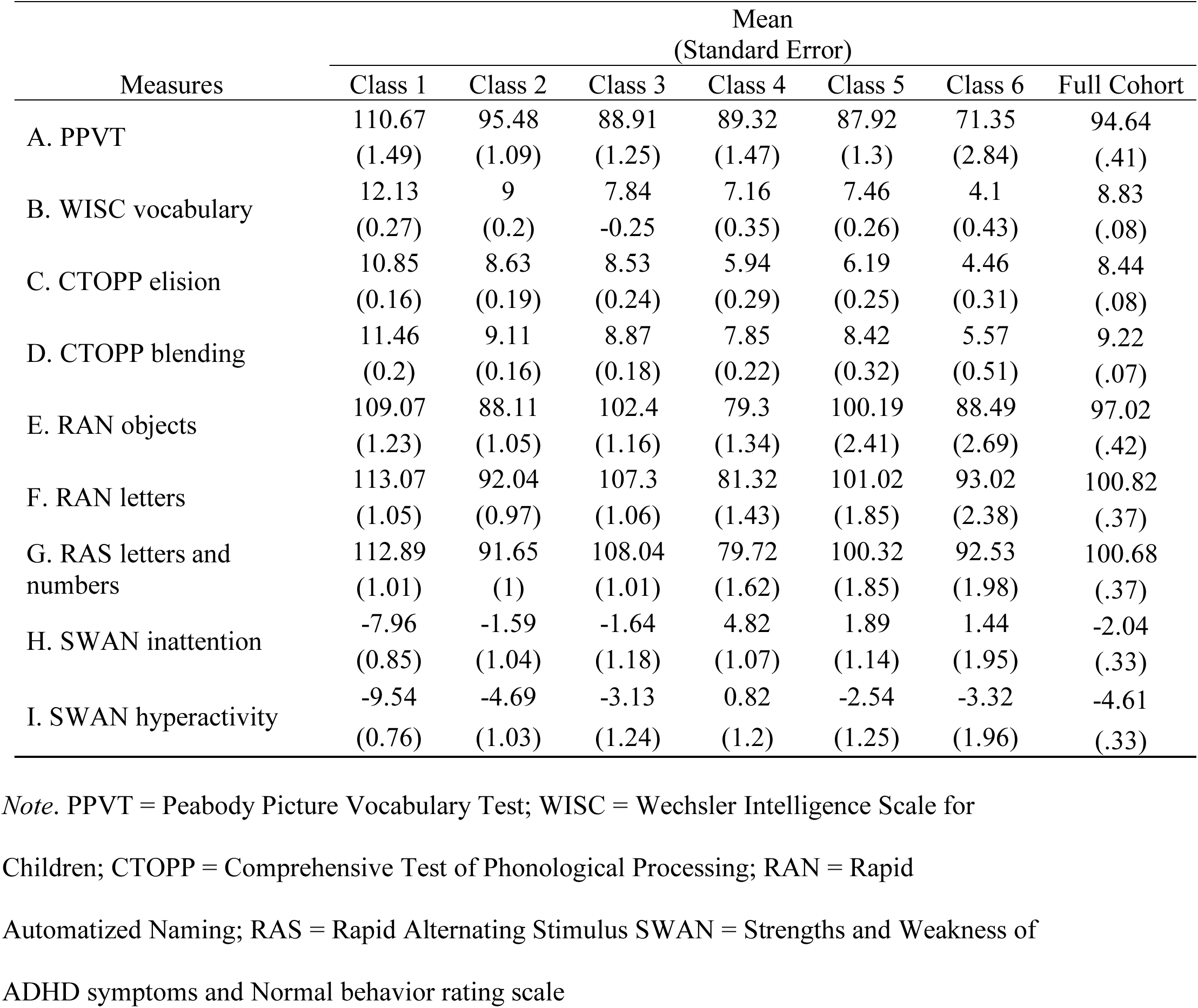
Estimated means and standard errors for six groups on reading-related measures

